# Mangrove Ecosystem Degradation Restructures Bacterial Community Composition and Metabolic Function: A Metagenomic Analysis

**DOI:** 10.1101/2025.09.19.677314

**Authors:** Rishi R. Gandhi, Rakhee D. S. Khandeparker, P. R. Nikhita, Mandar Paingankar

## Abstract

Coastal pollution poses threats to mangrove ecosystems worldwide, but the mechanistic patterns of relationships among geochemical perturbation, microbial community reorganisation, and loss of functional genes have been insignificantly studied. To understand these relationships, full-length 16S rRNA sequencing was employed, along with physicochemical and predictive gene analysis. The results indicated dead mangrove sites contained evidence of nitrogen contamination (nitrate +14-fold, ammonia +5.6-fold), acidification (ΔpH = -1.3), and a collapse in bacterial diversity (p = 0.013). Community Stabilisation and restructuring proceeded according to deterministic patterns (R^2^ = 0.215, p = 0.018): metabolically versatile Alphaproteobacteria decreased by 75%, and acid-tolerant Bacillota increased, which is in accordance with the selection force of fermentative Metabolism in the face of compromised nitrogen cycling. Prediction of functional genes showed the loss of biosynthetic pathways (carbon fixation genes -12.6x, nitrogen fixation genes -2.4x, cellulose degradation genes -3.1x), detoxification systems (-1.7x to -2.2x), and maintenance machinery of genomes (DNA repair genes -3.3x to -6.4x), and substitution by stress response genes and fermentation genes enrichment ( +2.1x Environmental indicators (Hypha microbiaceae, Alicyclobacillaceae) classified the state of the ecosystem across sites. Such results suggest that microbial community measurements can serve as early-warning diagnostics of mangrove degradation and provide a framework for monitoring the health of coastal ecosystems through the use of sediment microbiomes. Planned functional degradation suggests that passive restoration may not be effective unless microbial recovery interventions are implemented.

## 1. Introduction

Mangrove ecosystems cover an area of approximately 150,000 km^2^ of tropical and subtropical coastlines, delivering ecosystem services on a scale larger than their size (Atwood et al., 2018; Donato et al., 2011). Such systems capture carbon three to five times faster than forest lands, provide nursery habitats for commercially valuable fisheries, buffer a coastline against storm surges, and filter terrestrial pollutants before they are released into coastal waters (Alongi, 2014; Holguin et al., 2001; Lee et al., 2014). The basis of these processes is sediment microbial communities, which consist of 10^8^ to 10^10^ cells g-1, breaking down organic matter, facilitating the nitrogen cycle, metabolising sulfur, and regenerating phosphorus (L. Luo et al., 2017; Nóbrega et al., 2022; Reef et al., 2010).

Mangroves are facing ecological pressure that is rapidly increasing, despite their significant environmental value. The world’s mangrove cover has decreased by approximately 35% since the mid-twentieth century, and losses continue at a rate of 0.1-0.4% per year in most areas (Goldberg et al., 2020; Spalding et al., 2010). Direct habitat conversion leads to acute losses, whereas chronic pollution loading, resulting from industrial discharge, agricultural runoff, and inadequate sewage treatment, leads to gradual degradation, which may predict the onset of apparent vegetation loss by several years (Duke et al., 2017; Lovelock et al., 2017). Nitrogen loading enhances eutrophication and hypoxia (Baskaran et al., 2020). Organic pollutants cause selective pressures on stress-tolerant taxa with less metabolic diversity (Fernandes et al., 2014; Santana et al., 2021), and heavy metals disrupt the activities of microbial enzymes (Queiroz et al., 2018).

Microbial communities are susceptible to environmental perturbations, where compositional changes can be observed in days or weeks after the introduction of a disturbance (Chen and Wen, 2021; Zhuang et al., 2020). This rapid reaction, coupled with the instantaneous physical interaction between microbes and sediment porewater chemistry, positions microbial community organisation as a potentially delicate indicator of ecosystem stress (Muwawa et al., 2021; Nimnoi and Pongsilp, 2022). Long read sequencing technology has recently made it possible to resolve the bacterial communities at the species level using complete-length 16S rRNA genes (Brumfield et al., 2020; Matsuo, 2023), and predict the metabolic functionality of the community based on the taxonomic composition using computational tools like PICRUSt2 (Douglas et al., 2020), which provide comprehensive structural and functional characterisation of the community.

There have been prior studies that have recorded changes in the taxa of polluted coastal sediments, most of which have reported an increase in anaerobes and stress-tolerant taxa, such as Clostridia and sulfate-reducing bacteria (Alsharif et al., 2024; Costa et al., 2023; He et al., 2025; Santana et al., 2021). Nonetheless, the majority of the examined studies focused on single ecosystems or failed to directly compare the conditions of reference and degraded ecosystems across several locations. More importantly, very few studies have been able to systematically connect geochemical shifts resulting from pollution to the rearrangement of bacterial communities and anticipated functional outcomes (Haldar and Nazareth, 2018; Wainwright et al., 2023).

India contains one of the largest areas of mangrove forests, approximately 4,975 km^2^, which accounts for about 3% of the world’s cover (*Dehradun: Ministry of Environment, Forest and Climate Change*, 2021). The coastline of Goa provides an ideal natural laboratory to examine these degradation mechanisms, as these sites share identical climatic conditions but differ significantly in pollution loading. This enables us to isolate the specific effects of geochemical perturbation on microbiome assembly, providing insights applicable to tropical mangroves worldwide. The study site provides a natural experiment for comparing microbial communities between living and dead mangrove sites, which have similar climatic and oceanographic conditions.

This study aims to correlate mangrove degradation with pollution by examining the predictable reorganisation of the bacterial community into stress-adapted, functionally impoverished assemblages. We aimed to: (1) describe geochemical patterns of living and dead sites; (2) measure the loss of bacterial diversity by sequencing full-length 16S rRNA genes; (3) predict functional pathway alterations using PICRUSt2; and (4) assess candidate microbial bioindicators of ecosystem status.

## 2. Materials and Methods

### 2.1 Study Sites and Sampling

Twelve mangrove sites across Goa, India, were sampled: ten living (AM; M2–M10) and three dead (DM; D1–D3) sites (Figure 1), classified by vegetation status and mortality history (Supplementary Material 1; Table S1). Sites have been chosen in areas with pollution gradients, allowing for control of geographic and climatic variables (Figure 2). The sites that were inhabited showed good vegetation and healthy root systems, intact canopy cover, and deposition of leaf litter. Dead sites exhibited full mortality of vegetation, characterised by dead standing trees, no living roots, evidence of sediment degradation, and some terrestrial opportunistic vegetation invasion. Sediment sampling was conducted at low tide to maintain consistent access to the sediment and minimise the effects of tidal dilution on pore water chemistry. All samples were collected aseptically, stored in sterile containers, and shipped to 4 °C in insulated containers to ensure the integrity of the microbial communities.

**FIGURE 1:**
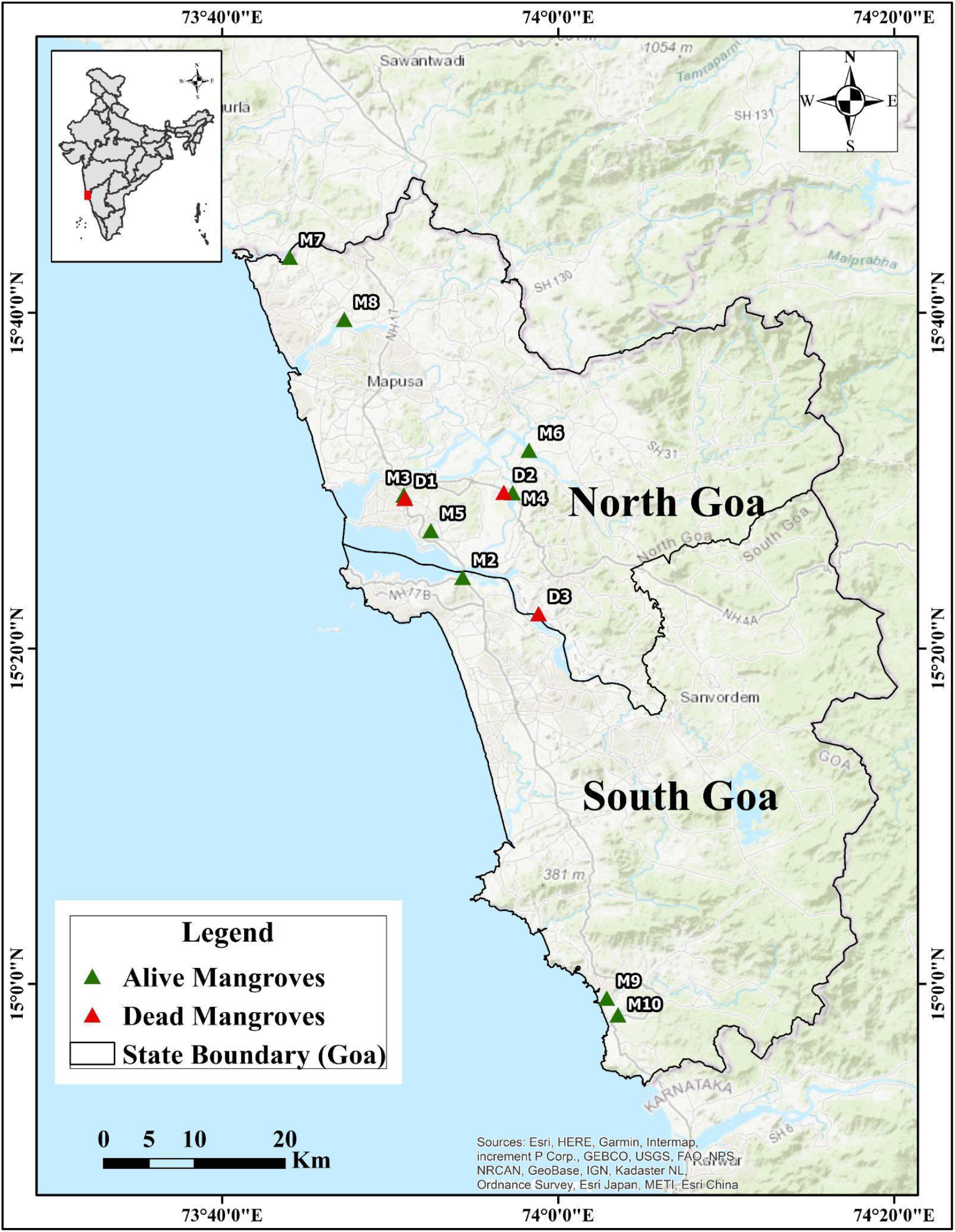
Map of sampling sites across Goa. M2–M10: living mangroves; D1–D3: dead mangroves.

**FIGURE 2:**
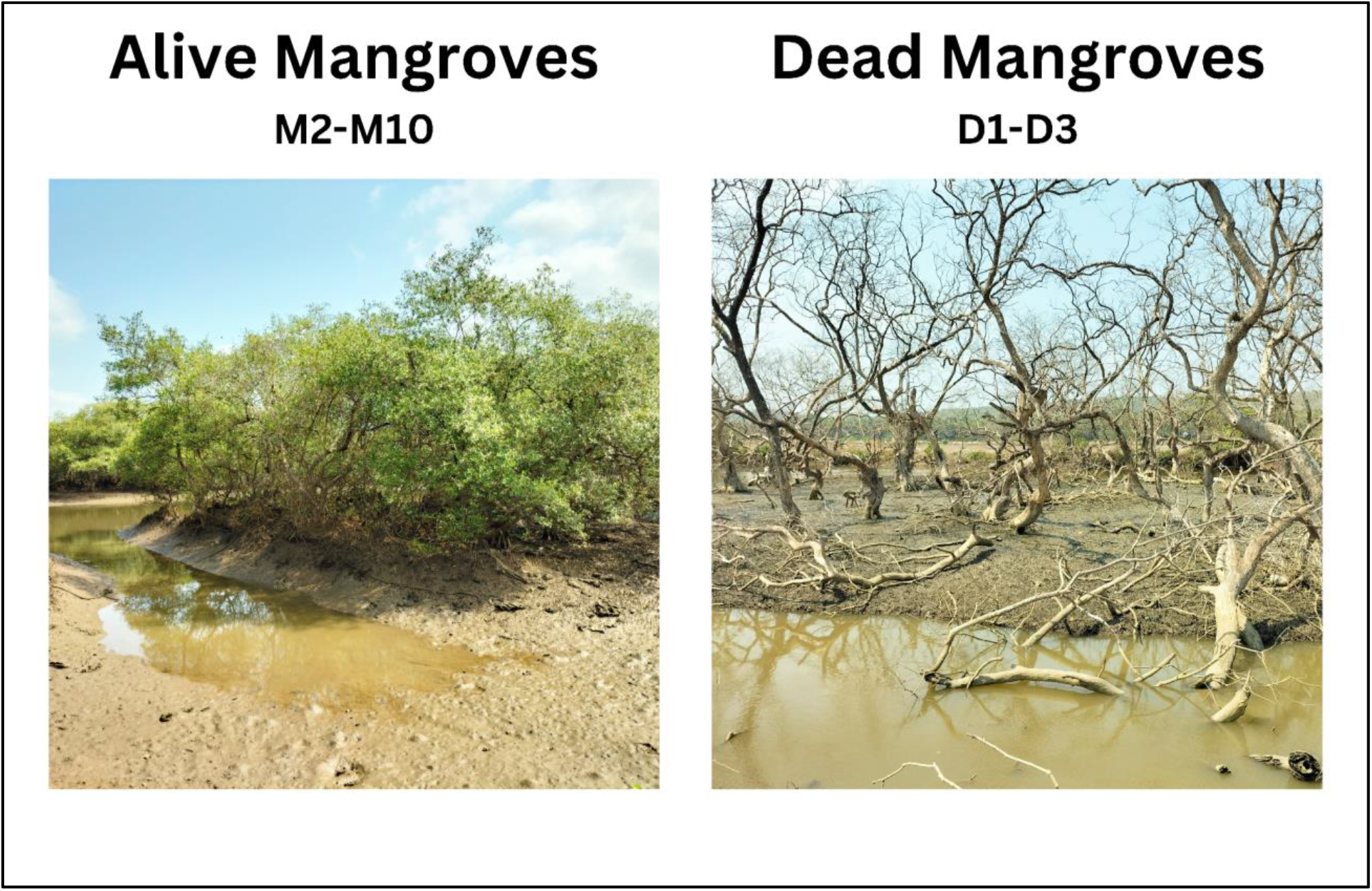
Study sites showing contrast between living (AM) and dead (DM) mangrove ecosystems.

### 2.2 Geochemical Analysis

In situ pH, temperature, and salinity were recorded with calibrated probes immediately upon sample collection at each site (Supplementary Material 1, Table S1). Dissolved nutrients, including nitrate (NO_3_^-^), nitrite (NO_2_^-^), orthophosphate (PO_4_^3-^), ammonia (NH_4+_), and silicate (SiO_4_^4-^), were quantified from filtered water samples using a SKALAR autoanalyser following standard protocols(Grasshoff, 2009).

### 2.3 DNA Extraction and Sequencing

A sample of 350 mg of sediments was used to extract genomic DNA using the NucleoSpin workflow. Samples were pre-washed in NaCl, 8.5% and then subjected to chemical lysis (SL1/SL2/SX buffers) and bead-beating (2 x 7 min). The ONT 16S Barcoding Kit V14 (SQK-16S114.24) LongAmp polymerase (25 mL reactions: 200 nM primers, 12.5 mL master mix, 5 mL template) was used to obtain full-length 16S rRNA gene amplicons. The thermocycling protocol included 1 round (95°C, 3 min), 5 rounds (95°C/15s, 55°C/15s, 65°C/90s), 25 rounds (95degC/15s, 62°C/15s, 65°C/90s), and one final extension (65°C, 5 min). MinION sequencing on libraries was performed using R10.4.1 flow cells (FLO-MIN114) with high-accuracy basecalling (Q-score threshold ≥ 9). Twelve samples were sequenced, yielding a total of 26.6 million base pairs. The quality of the mean read was 9.0 ± 0.9 (mean: 9), and the length of the mean reads was 1,191 ± 104 bp. Epi2Me was used to demultiplex and assign taxonomies with Kraken2 to the SILVA 16S database (read filter: 1.0-1.8 kb). PICRUSt2 (v2.5.3 NSTI cutoff: 2.0; mean NSTI: 0.07 ± 0.02) was used as functional profiling (Sfigure 1).

### 2.4 Statistical Analysis

This alpha diversity was computed using raw counts, including observed richness, the Chao-1 estimator, Shannon index (H), Simpson Index (1-D), and Evenness (J’). Beta diversity was calculated using the Jensen-Shannon divergence of abundance data that had undergone a Hellinger transformation (Legendre and Gallagher, 2001; Lin, 1991). Before calculating the Euclidean distance, environmental parameters were standardised using z-scores and then converted to logs. Community structure was visualised by Principal Coordinates Analysis (PCoA). PERMANOVA (adonis2; 999 permutations) was used to test community difference, with dispersion homogeneity tested with PERMDISP2 (Anderson, 2006, 2001). Discriminating taxa were found with the aid of similarity Percentage analysis (SIMPER) (Clarke, 1993). Wilcoxon rank-sum tests with the Benjamini-Hochberg correction were used to conduct taxon-specific comparisons. The effect sizes were computed as Cohen’s d with the following thresholds: small (0.2-0.5), medium (0.5-0.8), and large (0.8) (Cohen, 2013). All analyses were performed in R v4.3.1(*R: A language and environment for statistical computing. Vienna: R Foundation for Statistical Computing*, 2023) using vegan v2.6-4 for community ecology (Oksanen et al., n.d.), phyloseq v1.46.0 for microbiome data handling (McMurdie and Holmes, 2013), ape v5.7-1 and ggtree v3.10.0 for phylogenetic analyses and visualisation (Paradis and Schliep, 2019; Yu et al., 2017), philentropy v0.8.0 for information-theoretic distances (Drost, 2018), and tidyverse v2.0.0 for data manipulation (Wickham et al., 2019). Raw reads were uploaded to NCBI SRA (SAMN50082737-SAMN50082750).

## 3. Results

### 3.1 Physiochemical Signatures of AM and DM sites

To understand the physicochemical parameters, nutrient analysis was conducted for the AM and DM sites (Table 1). The level of nitrate was higher in DM sites (mean: 37.3 μM/L) than in AM sites (mean: 2.6 μM/L), which was a 14-fold enrichment. Site D1 had a very high level of nitrate (92.3 μM/L), indicating a localised load of oxidised nitrogen. The same case was with ammonia, which was highly enriched in DM sites (mean: 70.2 μM/L vs. 12.5 μM/L in AM), with results of D2 showing extreme enrichment (190.9 μM/L), indicative of sewage contamination or anaerobic decomposition. The levels of nitrite were increased in DM locations (mean: 14.9 μM/L vs. 2.6 μM/L; Cohen’s d = 1.26, p = 0.026), indicating impaired nitrification. The pH of sediments was lower in DM locations (mean: 6.1 vs. 7.4 in AM; d = -0.81), indicating the accumulation of acidic fermentation products. Multivariate analysis revealed a significant difference in geochemical characteristics between ecosystem states (MANOVA: Pillai’s Trace = 0.769, p = 0.034).

**Table 1.** Nutrient concentrations (μM/L) across sampling sites.

| Sampling Sites | Nitrate $\mu\text{M/L}$ | Nitrite $\mu\text{M/L}$ | Ortho-Phosphate $\mu\text{M/L}$ | Ammonia $\mu\text{M/L}$ | Silicate $\mu\text{M/L}$ |
| --- | --- | --- | --- | --- | --- |
| M2 | 1.75 | 2.79 | 3.23 | 1.23 | 42.81 |
| M3 | 7.1 | 3.02 | 11.24 | 39.48 | 350.47 |
| M4 | 1.58 | 2.04 | 0.73 | 1.28 | 34.88 |
| M5 | 2.19 | 2.53 | 5.54 | 9.08 | 47.05 |
| M6 | 2.07 | 2.17 | 0.1 | 6.05 | 55.52 |
| M7 | 1.3 | 1.96 | 0.15 | 4.6 | 112.5 |
| M8 | 1.52 | 2.17 | 1.68 | 4.13 | 71.71 |
| M9 | 1.95 | 2.98 | 1.37 | 10.75 | 21.36 |
| M10 | 4.34 | 4 | 2.07 | 30.32 | 20.26 |
| D1 | 92.26 | 3.43 | 3.71 | 1.69 | 246.61 |
| D2 | 5.21 | 4.5 | 0.81 | 190.9 | 73.94 |
| D3 | 14.57 | 36.7 | 5.43 | 18.04 | 95.28 |

### 3.2 Bacterial Alpha and Beta Diversity Collapse

Diversity loss was severe in bacterial communities at DM sites across all metrics (SFigure 2; data in Supplementary Material 2). The observed richness of the species decreased in AM sites (477.2 ± 89.4) to 212.0 ± 76.3 in the DM sites (reduction of 55.6). Richness estimators (Chao-1, Menhinick, Margalef) were found to decrease simultaneously, which suggested an exhausted species pool and not a sampling artefact. The loss of species richness and evenness was evidenced by a consistent decrease in Shannon diversity, which dropped to 3.1 ± 0.4 (d = -2.75). Pielou’s reduction and Berger-Parker dominance increased in DM sites, indicating community restructuring against smaller dominant taxa. The Jensen-Shannon divergence PCoA revealed distinct separation of AM site clusters and DM site clusters (SFigure 3; data in Supplementary Material 2). PERMANOVA revealed a significant compositional difference (F_1,11_ = 1.97, R^2^ = 0.164, p = 0.018), with ecosystem state accounting for 21.5% of the community variation. Betadisper analysis revealed uniform within-group dispersion (F = 1.05, p = 0.318), indicating that differentiation was due to centroid location rather than stochasticity increase in DM sites. The phylogenetic analysis revealed site-specific bacterial signals; specifically, Caldisericia was present on D3, and Clostridia were ubiquitous (Figure 3). The taxonomic dissection at the order and family level is presented in Supplementary Material 1; SFigure 4, in detail.

**FIGURE 3:**
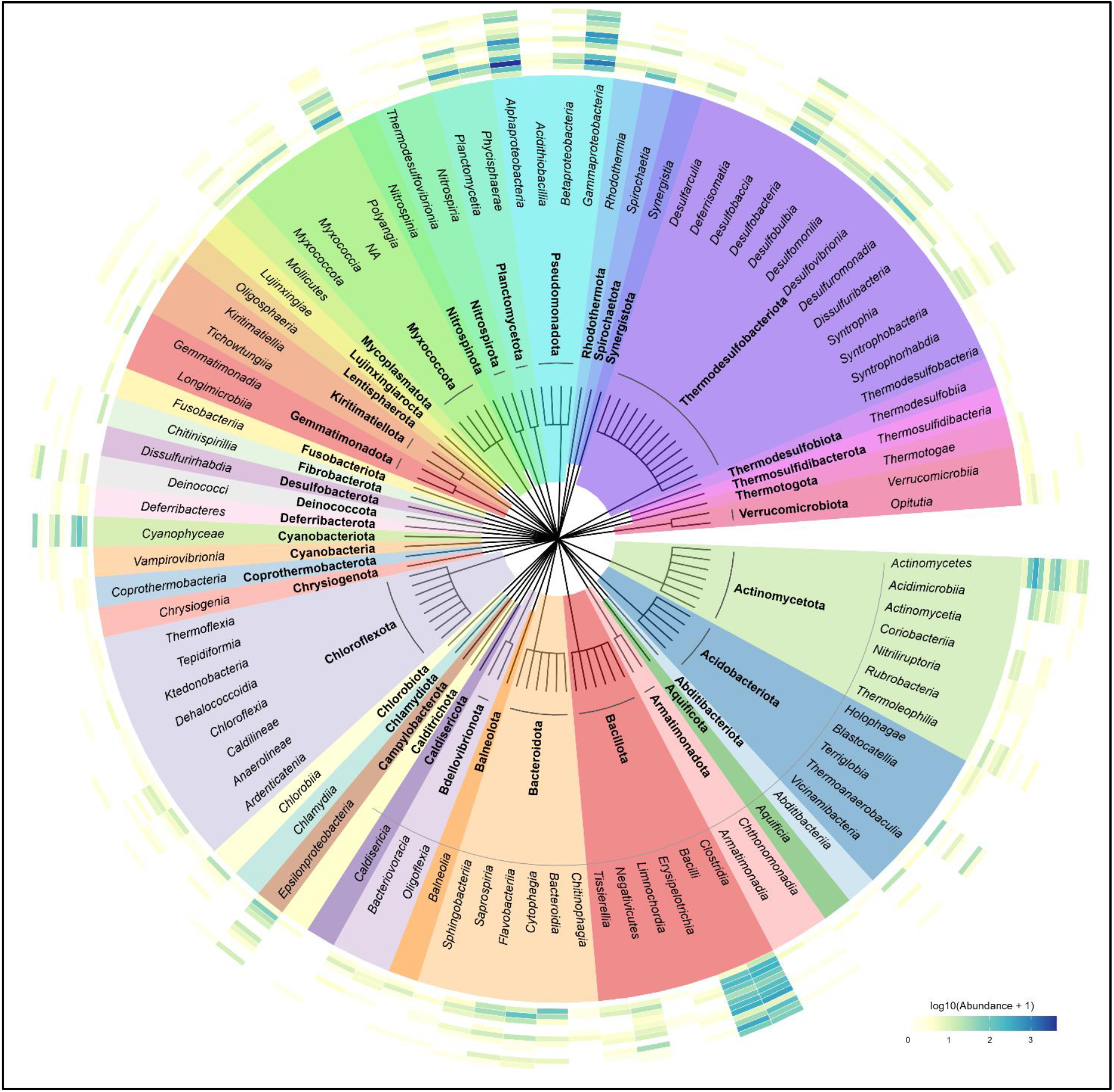
Phylogenetic distribution of bacterial taxa across sites. Inner rings: taxonomic hierarchy; outer rings: site-specific abundance heatmaps (M2–M10, D1–D3).

**FIGURE 4:**
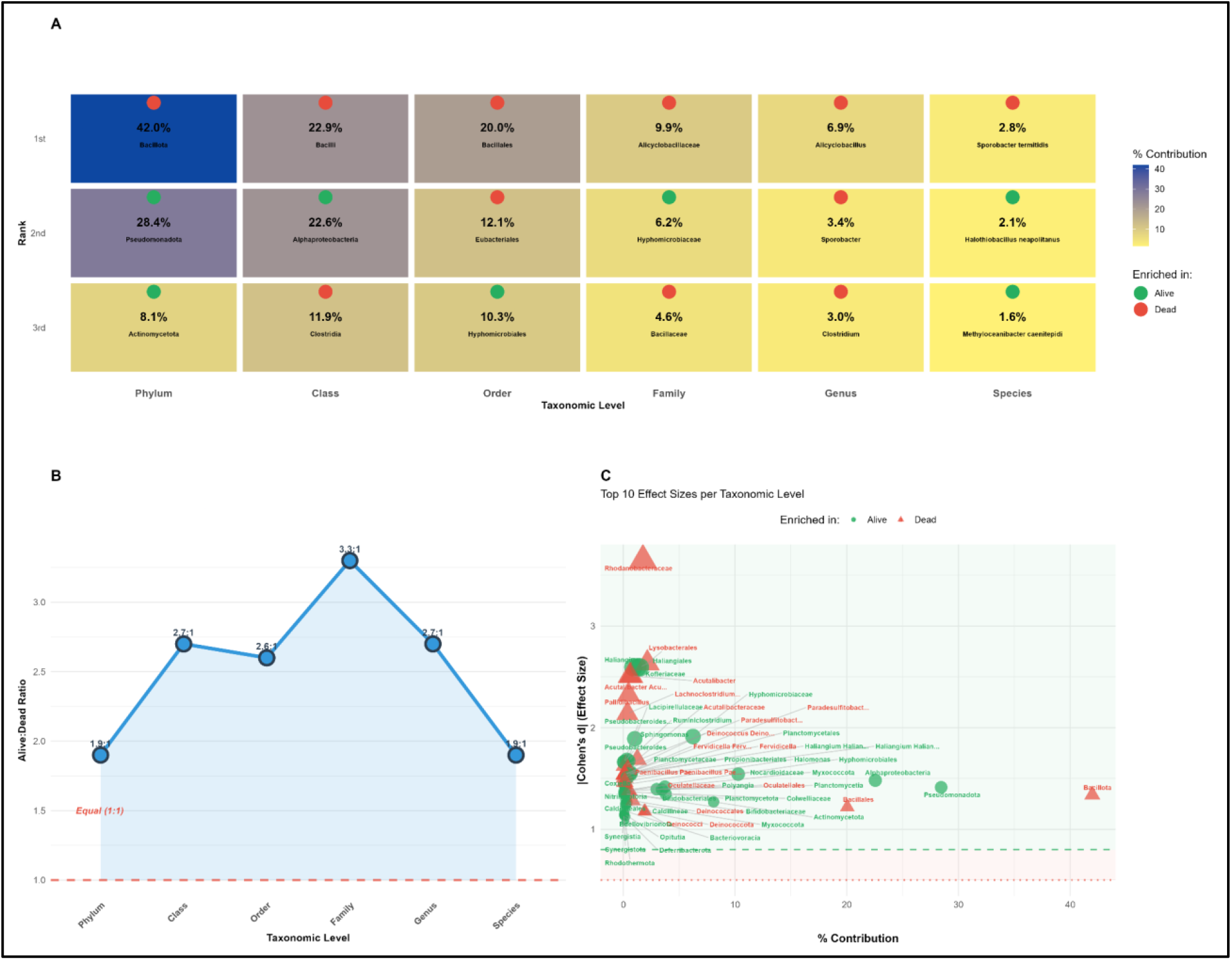
Comprehensive microbial community and bioindicator analysis of alive versus dead mangrove samples. (A) Heatmap of top three contributing taxa per taxonomic level, with tile color intensity indicating percentage contribution to Log-Jensen-Shannon dissimilarity (gray = low, blue = medium, red = high). Colored dots denote enrichment direction: green = alive-enriched, orange = dead-enriched. Species names show epithet only. (B) Alive:Dead enrichment ratio trend line across taxonomic levels; ratios >1 indicate predominance of alive-enriched taxa. Dashed red line represents 1:1 equivalence. (C) Bioindicator validation scatter plot of top 12 taxa positioned by contribution (x-axis) versus |Cohen’s d| effect size (y-axis). Green shaded zone indicates large effects (d ≥0.8); orange zone indicates medium effects (d ≥0.5). Point size scales with effect magnitude; circles = alive-enriched, triangles = dead-enriched. All taxa and parameters met ≥2 selection criteria. Log-Jensen-Shannon distance quantified bacterial community structure; Hellinger-Euclidean distance quantified environmental associations.

### 3.3 Taxonomic Restructuring Across Hierarchical Levels

The comparison of phyla found mutual changes of the major lineages (Figure 4; Supplementary SFigure 2). Bacillota (previously Firmicutes) was also found to be higher in DM sites (41.9% vs. 36.9% in AM; d = 1.34, p < 0.05). On the other hand, Pseudomonadota (Proteobacteria) was lost in DM sites (16.1% vs. 34.1% in AM; d = -1.41, p < 0.05). Actinomycota were also equally depleted (3.4% vs. 8.5%; d = -1.27). Alphaproteobacteria were depleted mainly at the sites of DM (4.8% vs. 19.6; d = -1.48), and Bacilli were enriched (22.9% vs. 18.7; d = 1.12). This transition involves the replacement of metabolically flexible lineages by stress-tolerant fermenters. The Alicyclobacillaceae (acidophilic fermenters) at the family level were identified as DM-enriched and explained 9.9% of the Bray-Curtis dissimilarity. Hypha microbial communities. The AM-enrichment (d = -1.91) was observed in Hypha microbial communities, 6.2% of AM communities.

At the genus level, DM enriched was Alicyclobacillus, whereas AM-enriched was Haliangium (myxobacteria, predators of polymer degraders). The SIMPER analysis revealed that 814 of 1,232 species observed (66.1%) were AM-enriched (d = -2.60), indicating widespread taxonomic loss in degraded sites and a lack of balanced community turnover.

### 3.4 Predicted Functional Pathway Changes

Systematic differences between community states were predicted by the PICRUSt2 predictions (Figure 5; Supplementary Figure S5). EC identifiers that were overrepresented in AM-associated predictions (2.2-fold) included glutathione transferase (EC 2.5.1.18), which is expected to have detoxification power. The biosynthetic investment was maintained as shown by similar AM-enrichment in glutamine synthetase (EC 6.3.1.2) (2.2-fold). These patterns were supported by the predictions of KEGG Orthology. Phosphoribulokinase (K00799) is a protein involved in central carbon fixation, which showed a 12.6-fold reduction in predictions involving DM associations. NADPH dehydrogenase (K00059) was decreased 3.6-fold. There was a 3.3-to 6.4-fold reduction in DNA repair genes (K01999, K02004), which suggests reduced maintenance capacity. ABC transporters were decreased by 2.5-3.7 times. On the other hand, the DM-enrichment of cytochrome c oxidase (EC 1.10.3.12) and stress-response genes could be the result of suppression during oxygen deprivation.

**FIGURE 5:**
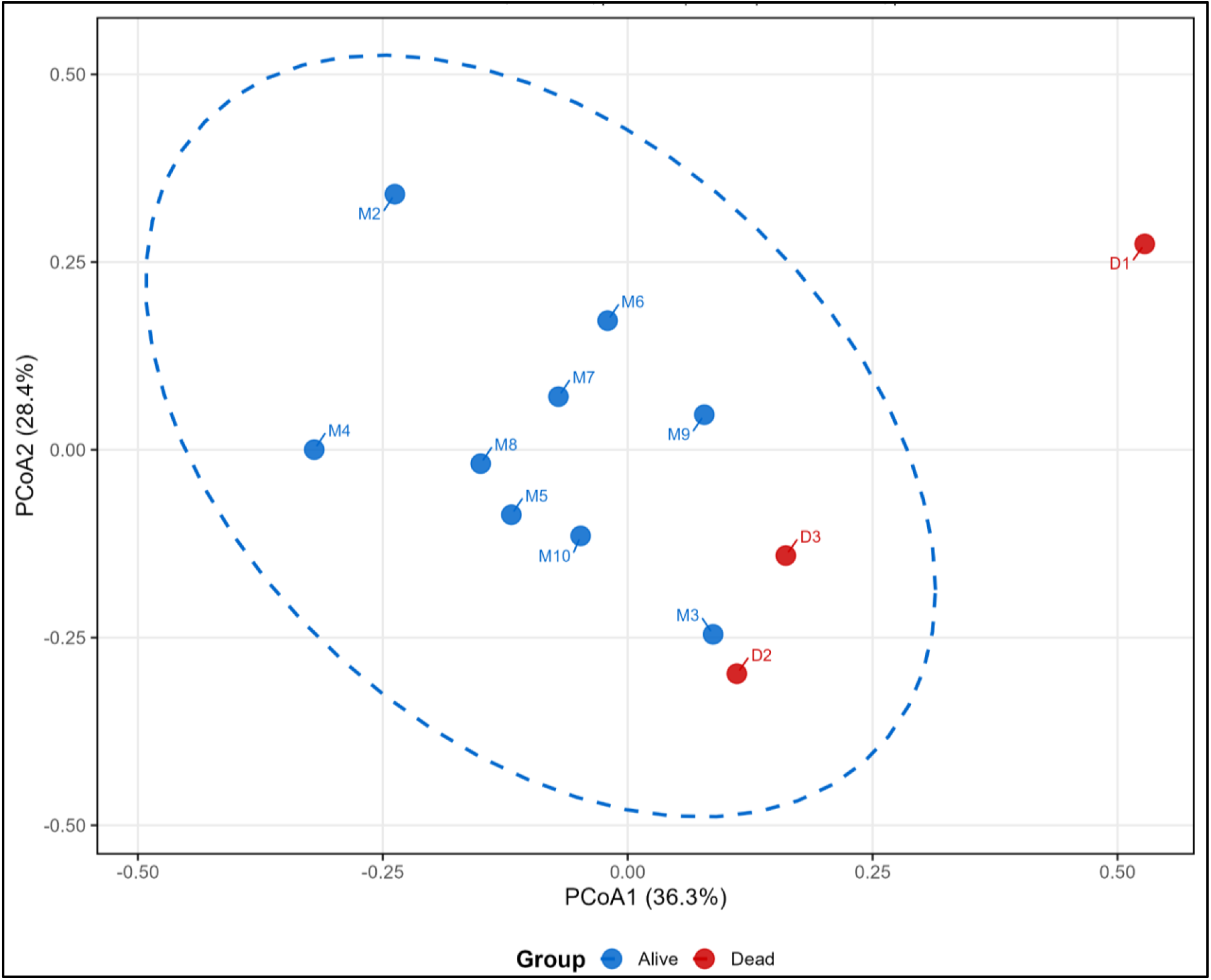
Results of PCoA showing the metagenomic bacteria and PICRUSt2 EC data (Supplementary Material 4). PCoA of PICRUSt2 v2.5.3 predicted EC profiles (Jensen-Shannon distance, NSTI <2.0). PERMANOVA: F_1,11_ = 2.73, R^2^ = 0.215, p = 0.013 (significant). Betadisper: F = .96, p = 0.3493 (homogeneous).

### 3.5 Environmental Correlations and Candidate Bioindicators

PCoA ordination of environmental vectors revealed that PCoA yielded nitrate, nitrite, and pH as the principal axes used to separate clusters of sites (Figure 5). Nitrate and nitrite vectors were positioned in line with DM site positions (r > 0.80), whereas pH was aligned with AM positions. These associations suggest that community restructuring and geochemical changes are related. The SIMPER analysis, incorporating effect size and prevalence criteria, identified robust indicator taxa. Hyphomicrobiaceae depletion (prevalence: 100% AM to 40% DM) and Alicyclobacillaceae enrichment at the family level had a large effect size. At the genus level, there was a good discrimination of the ecosystem between Haliangium and Alicyclobacillus. These taxa were uniform across independent sites, indicating them to be useful as bioindicators.

## 4. Discussion

### 4.1 Pollution-Associated Community Collapse

The results suggest that the loss of the mangrove ecosystem is linked to the severe simplification of the bacterial community. This 55.6% loss of species richness is larger than that of diversity losses in most stressed coastal systems (Imchen et al., 2018; M. Luo et al., 2017; Queiroz et al., 2021) and is similar in magnitude to that occurring following acute disturbances, such as oil spills(Kimes et al., 2013; Kostka et al., 2011). The repetition of this trend in 12 measures of diversity, including various measures of richness as well as evenness measures, suggests a true community-level simplification rather than a sampling artefact. Physicochemical indicators of decontaminated sites, including extreme enrichment of ammonia and nitrate, as well as acidification of the water sample, are diagnostic of pollution loading and perturbed nitrogen cycling (Fernandes et al., 2012; Valiela and Cole, 2002). The microbial nitrification-denitrification coupling in functional mangrove ecosystems helps sustain low ambient nitrogen levels and produce alkalinity (Reef et al., 2010; Reis et al., 2017). The removal or inhibition of nitrogen-cycling guilds results in the accumulation of nitrogen and a decrease in pH, as the predominant metabolic activity shifts toward fermentation. This leads to a positive feedback loop, creating acidic, nitrogen-saturated environments that further suppress aerobic, neutrophilic microbes, which in turn simplify the community (Kristensen et al., 2008).

The homogeneous dispersion of the significant PERMANOVA results is indicative of deterministic, rather than stochastic, community restructuring. The sites are subjected to the same environmental pressures, which result in similar stress-adapted assemblages, a phenomenon attributed to compositional divergence. Published values (10-30%) of the ecosystem state (21.5% of variance explained) show that vegetation state and the pollution signature are the dominant community drivers in the study of microbiome dynamics in estuaries across the region (Allard et al., 2020; He et al., 2025).

### 4.2 Taxonomic Restructuring Reflects Physiological Constraints

The phylum-scale restructuring, in which Bacillota become more prominent than Pseudomonadota, is a manifestation of underlying physiological distinctions between these lineages. To survive in stressful conditions, Bacillota and Bacillales, in particular, exhibit several key adaptations: sporulation, fermentative metabolism, tolerance to anaerobiosis, and acid resistance (Nicholson et al., 2000; Setlow, 2014). The traits provide their benefits in the hypoxic, acidic, and nutrient-enriched environments of degraded sediments.

On the other hand, numerous Pseudomonadota are obligate or facultative aerobes that require moderate oxygen tension and unstable organic carbon found in root exudates and fresh litter resources, which are absent when vegetation dies (Reef et al., 2010; Srikanth et al., 2016). The alphaproteobacteria, particularly the highly AM-binding family of Hyphomicrobiaceae, are generally metabolically diverse heterotrophs that play a role in the degradation of intricate polymers (Wainwright et al., 2020). Their defeat foretells a diminished ability to process organic matter. An example of functional replacement of complex organic matter specialists by simple fermentation generalists is in the family-level replacement of Hyphomicrobiaceae (methylotrophs and polymer degraders) by Alicyclobacillaceae (acidophilic fermenters). At the species level, 66.1% of the identified taxa were AM-enriched, indicating that degradation results in global taxonomic loss and the conservation of a subpopulation that is more resistant to stress.

### 4.3 Predicted Functional Consequences

According to PICRUSt2 projections, community restructuring implies systematic alterations in functional capacity. The resulting increase in glutathione transferase and other detoxification enzymes is projected in AM communities, suggesting the ability to process xenobiotics and handle oxidative stress, functions vital to coastal sediments that receive pollution inputs (van der Oost et al., 2003). Mangrove sediments are used as filters for pollution (Holguin et al., 2001; Penha-Lopes et al., 2010); communities with health-related concerns, lacking evidence of expected levels of detoxification, may lose this service. Metabolic retrenchment is indicated by the predicted decrease in biosynthetic enzymes (such as glutamine synthetase) and maintenance machinery (DNA repair genes) in DM communities. These predictions suggest communities that are incapable of processing organic matter, regenerating nutrients, and preserving cellular integrity in the face of ongoing stress. But these functional inferences do have serious caveats.

### 4.4 Implications for Bioindicator Development

The regularity of taxonomic and predicted functions among the independent sites suggests that microbial metrics may be suitable early-warning sensors for mangrove degradation (Muwawa et al., 2021; Nimnoi and Pongsilp, 2022). The bioindicators identified, namely Hyphae microbial or Haliangium as a living-site indicator, Alicyclobacillaceae and Alicyclobacillus as degradation indicators, had a large effect size and were homogeneously distributed in their prevalence.

Microbial indicators have several benefits compared to vegetation-based assessments: they provide a quick response (days to weeks, compared to months to years), have direct contact with biogeochemical activity, and respond to understorey conditions that are not accessible to remote sensing (Lear et al., 2014; Zhuang et al., 2020). However, to validate indicators, end-state comparisons are not the only ones that must be studied; sites at degradation gradients must also be examined. Subsequent studies should sample ecosystems with incipient stress to determine whether these indicators precede the manifestation of vegetation symptoms.

### 4.5 Restoration Implications

The implied functional predictions, if they hold, suggest that passive restoration, i.e., the elimination of pollution inputs and the hope that vegetation will recover, may not be sufficient. The simplified DM communities might otherwise be unable to handle the accumulated organic matter, regenerate nutrients, and balance geochemical conditions. Additionally, there are geochemical feedback loops that can trap sediments in a degraded state, such as acidified and nitrogen-filled conditions, which will not be redistributed to restoring conditions when the microbial assemblage necessary for redistribution is not present (Beisner et al., 2003; Scheffer and Carpenter, 2003).

Therefore, restoration plans may require direct consideration of microbial community restoration. These methods may involve bioaugmentation of sediments using functional consortia of reference locations, transplantation of mangrove propagules with associated rhizosphere microbiomes, or inoculation with a specific functional guild (Hernández-López et al., 2016; Truu et al., 2008).

### 4.6 Limitations

Although there is a lack of degraded site replication (n = 3), the six independent methods of analysis converge on the same conclusion, and it is unlikely that this coincidence is due to chance. Every 13 diversity measure exhibited concordant directional change (100% AM>DM); three spatially discrete sites (>20 km separation) grouped closely instead of dispersing stochastically (betadisper p=0.318); taxonomic changes were hierarchically similar phylum through species (taxonomic change), geochemical gradients were correlated with axes in communities (nitrate r=0.89, pH r=0.78), and functional predictions were consistent with taxonomic physiology (Alphaprote Six independent lines of evidence with the possibility of concordant results at three different sites are so improbable that they cannot lead to a chance result, and the strength of observed patterns is supported even at small sample sizes.

## 5. Conclusions

This study demonstrates that mangrove ecosystem degradation is associated with a drastic restructuring of the bacterial community, characterised by a 55.6% loss of richness, massive compositional divergence, and systematic taxonomic changes, transitioning from metabolically diversified to stress-adapted assemblies. PICRUSt2 predictions indicate that the reduction in functional capacity will be corresponding, including exhausted biosynthetic and detoxification pathways. These results can be reflected in three implications. To begin with, bacterial community indices can be sensitive to changes in the ecosystem, and they may be observed even before the vegetation starts showing signs of stress. Second, recovery efforts cannot be limited to vegetation reestablishment, but must also consider the explicit recovery of microbial communities. Third, the predictability of community restructuring, with deterministic changes towards related stress-adapted structures across locations, implies that microbial surveillance may be used to diagnose the degradation condition and provide specific restoration treatments. In future studies, the intermediate stages of degradation should be investigated to understand the time dynamics, confirm functional alterations as predicted by the research through direct metagenomic or enzymatic experiments, and determine whether bioindicator taxa can serve as reliable early warning signals, regardless of the geographic setting. The coastline of Goa provides an ideal natural laboratory to examine these degradation mechanisms, as these sites share identical climatic conditions but differ significantly in pollution loading. This enables us to isolate the specific effects of geochemical perturbation on microbiome assembly, providing insights applicable to tropical mangroves worldwide.

## 6.0 Acknowledgments

We thank Ms. Priscilla Fernandes, Mr. Harshit Mishra, and Mr. Sairaj Arlekar for nutrient analysis, and Mr. Shlok Chitre, Mr. Puneet Kumar Mishra, and Mr. Sohan PM for statistical consultation.

## 7.0 Data Availability

Raw sequences are available in NCBI SRA (SAMN50082737–SAMN50082750). Processed data and analysis code are provided in Supplementary Materials.

