## Supplementary material 1 for "Mangrove Ecosystem Degradation Restructures Bacterial Community Composition and Metabolic Function: A Metagenomic Analysis": Supplementary material 1.docx

Supplementary table. 1: The table shows the pH, temperature, salinity, location and number of isolates from different sampling sites.

| **Site name** | **pH** | **Temperature**  **(℃)** | **Salinity (psu)** | **Location** |
| --- | --- | --- | --- | --- |
| **M2** | 7.6 | 32 | 40 | 15.403974545745497, 73.90517090806742 |
| **M3** | 7.2 | 28 | 65 | 15.485727914738789, 73.84680663690527 |
| **M4** | 7.4 | 32 | 35 | 15.487946895231579, 73.95488970806926 |
| **M5** | 7.7 | 32 | 40 | 15.450394197258788, 73.87357768619154 |
| **M6** | 7.1 | 30.5 | 29 | 15.530203833569303, 73.97102632341418 |
| **M7** | 7.1 | 29 | 25 | 15.721918852272518, 73.73357675496088 |
| **M8** | 7 | 30 | 36 | 15.660217751653718, 73.78767135476927 |
| **M9** | 7.9 | 35.5 | 40 | 14.985929276420753, 74.04818313325731 |
| **M10** | 7.6 | 33.5 | 35 | 14.969367229590466, 74.05930605376919 |
| **D1** | 7.3 | 31 | 10 | 15.482420201240176, 73.84780663690512 |
| **D2** | 3.5 | 38 | 49 | 15.488383291057886, 73.94604608108529 |
| **D3** | 7.5 | 32 | 39 | 15.367895406107504, 73.98051132905634 |

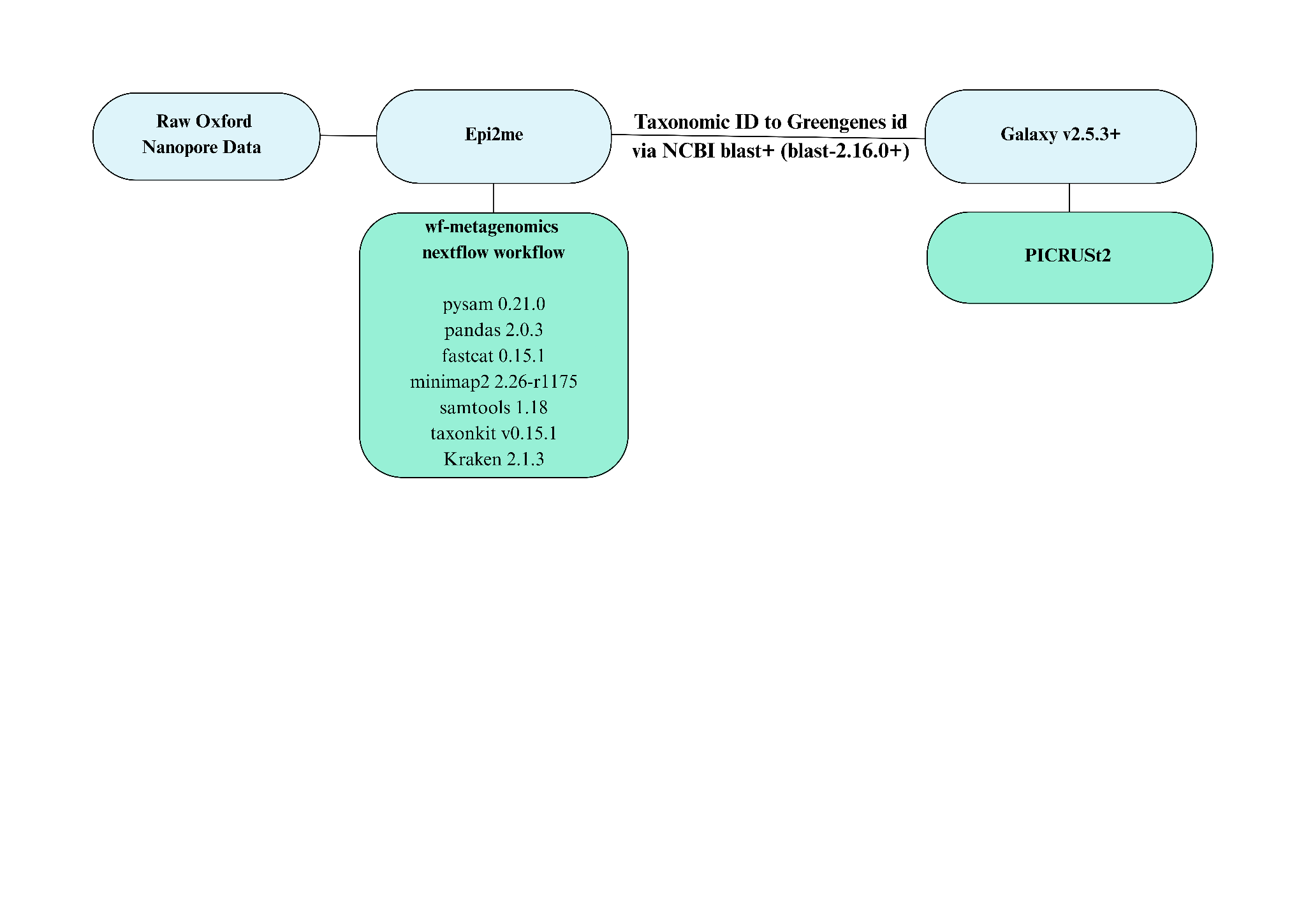

SFigure 1: Pipeline showcasing the metagenomic data processing steps. Oxford Nanopore (MinION, R10.4.1) reads processed via EPI2ME wf-metagenomics v2.11.0: quality control (fastcat v0.15.1, Q≥7, 1000-4000bp), taxonomic classification (minimap2 v2.26-r1175 → Kraken2 v2.1.3, SILVA 138 SSU, confidence 0.1), functional annotation (PICRUSt2 v2.5.3, BLAST+ v2.16.0+).

Following DNA sequencing using Oxford Nanopore technology, metagenomic analysis was performed using the EPI2ME wf-metagenomics Nextflow workflow (v2.11.0), which incorporates tools such as pysam (v0.21.0), pandas (v2.0.3), fastcat (v0.15.1), minimap2 (v2.26-r1175), samtools (v1.18), taxonkit (v0.15.1), and Kraken (v2.1.3). This pipeline generated two main output files: an abundance file containing bacterial names and their corresponding counts across sampling sites, and a taxonomic file with the corresponding taxonomic IDs. Since each file originated from a separate site, custom scripts were used to merge them into unified matrices that show bacterial distributions across all sites. For functional prediction, PICRUSt2 (v2.5.3+galaxy0) was used via the Galaxy platform, following the assignment of Greengenes IDs to taxa using BLAST+ (blast-2.16.0+, NCBI).

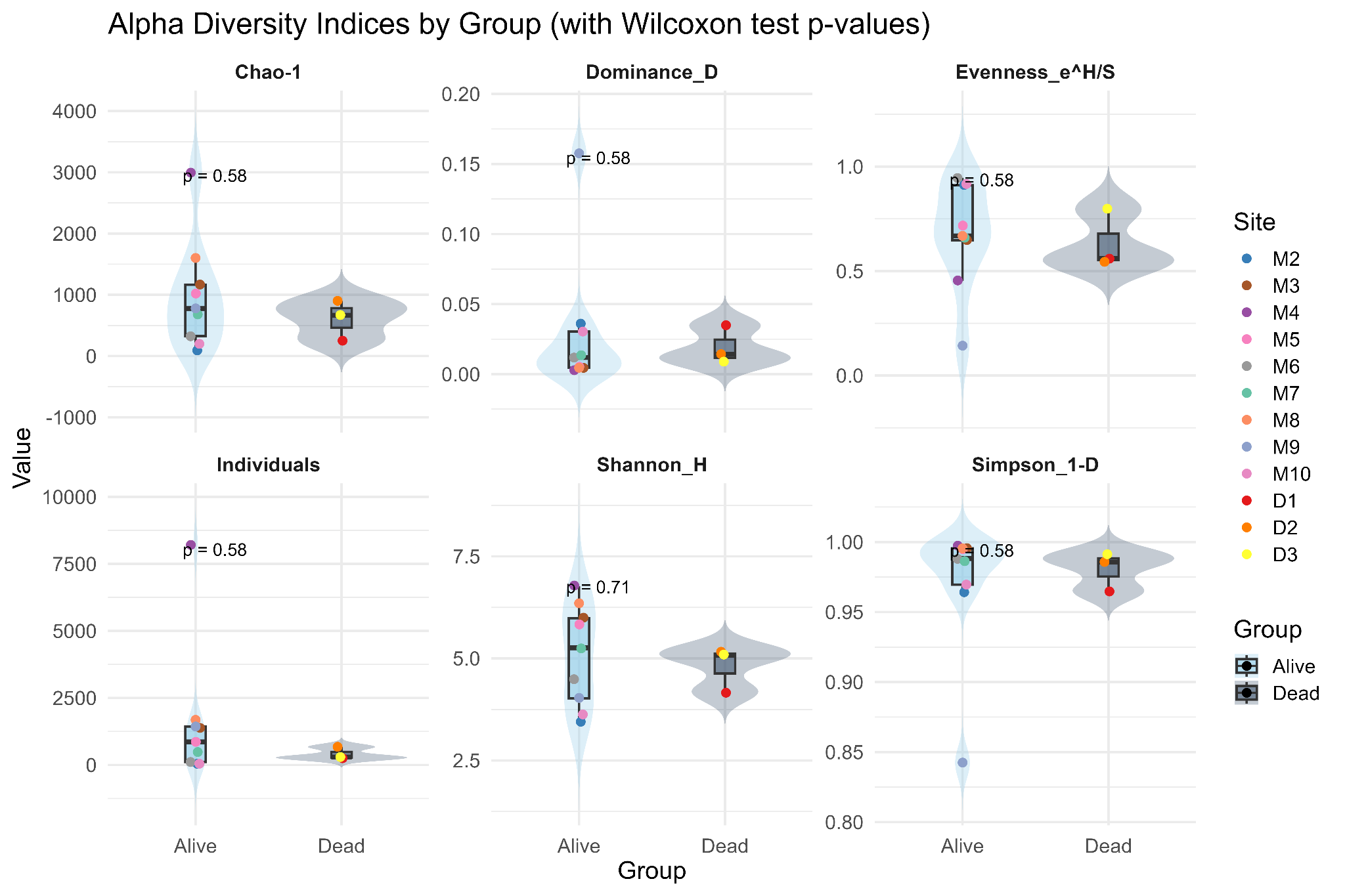

SFigure 2: Alpha diversity of the metagenomic genera. Diversity metrics (n=12 indices: Chao-1, Shannon_H, Simpson_1-D, Fisher_alpha, Dominance_D, Berger-Parker, Equitability_J, Evenness_e^H/S, Brillouin, Menhinick, Margalef, Taxa_S, Individuals) calculated using vegan v2.6-4 in R v4.3.1 for AM sites (n=9, blue) and DM sites (n=3, red).

Thirteen alpha diversity indices were calculated for each site, including observed species richness (Taxa_S, Individuals), dominance (Dominance_D, Berger-Parker), evenness (Equitability_J, Evenness_e^H/S), and diversity estimators (Shannon_H, Simpson_1-D, Fisher_alpha, Brillouin, Chao-1, Menhinick, Margalef). On a broader scale, species richness and diversity measures (e.g., Shannon-H, Simpson-1-D, and Chao-1) yielded higher values for the total sites that belonged to the AM group compared to the DM group. AM sites have always been more prosperous and diverse, with less dominance, whereas DM sites have been less affluent, with lower diversity indices and higher dominance, which means a loss of both species and an uneven distribution of relative abundance. The overall pattern of diversity measures towards community simplification in degraded sites shows a decrease in average richness of 55.6%.

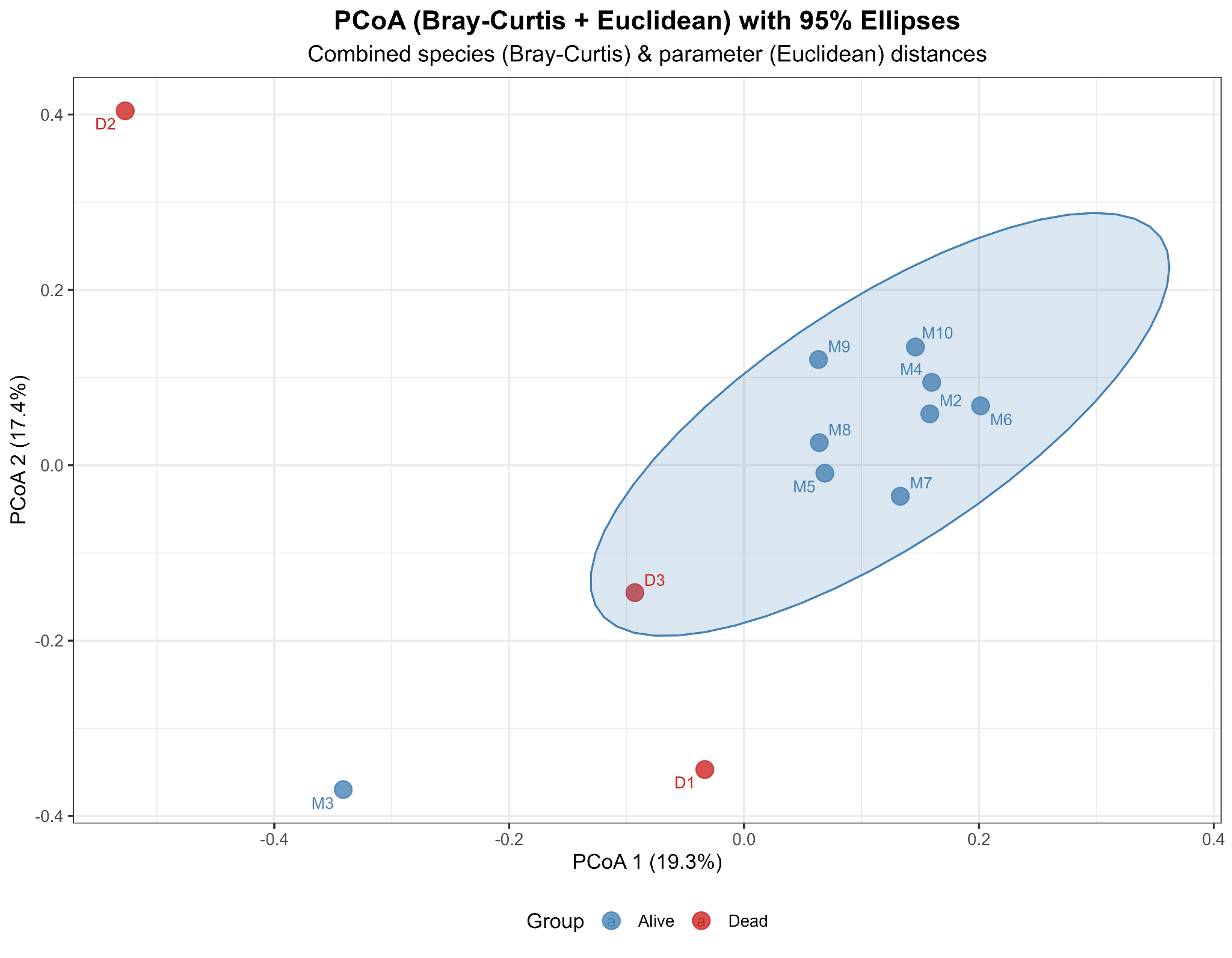

SFigure 3: PCoA showing the beta diversity of the metagenomic bacteria. Beta diversity (JSD-dissimilarity, Hellinger-transformed bacterial diversity and Euclidean, Log transformed environmental parameters data).

Principal Coordinates Analysis, based on Bray–Curtis dissimilarity (Hellinger-transformed bacterial counts) and log-Euclidean environmental parameters, revealed a clear separation between AM (blue) and DM (red) site clusters. PERMANOVA (F=1.97, *R*^2^=0.164, *p*=0.018) confirmed significant multivariate separation. Betadisper tests showed homogeneous within-group dispersion (F = 1.05, p = 0.318), indicating that the observed separation reflects true compositional divergence rather than differential variance. Dashed ellipses enclose 85–95% confidence regions for each group, emphasising minimal AM–DM overlap in ordination space.

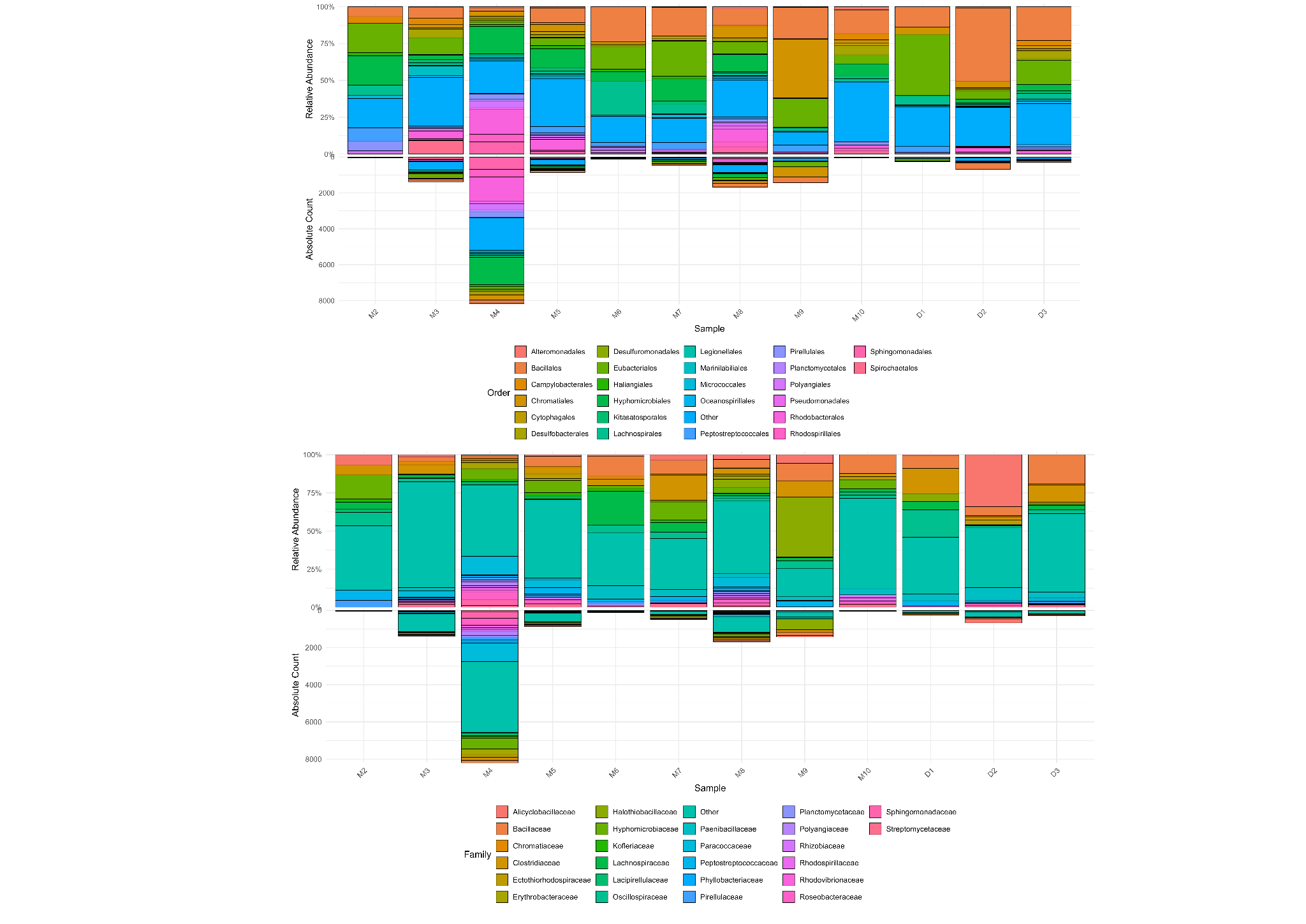

SFigure 4: Top 25 taxa of Order and family across all sampling sites. Relative abundance of top 25 taxa (Orders and Families) from 16S rRNA data.

Relative abundance of 16S rRNA of the top 25 taxa at Order and Family rankings across 10 alive mangrove (AM, blue) and 3 dead mangrove (DM, red) sites. There is apparent site-specific variation in dominant orders (Cytophagales, Sphingomonadales, Rhizobiales, Bacillales, Hyphomicrobiales) and families (Flavobacteriaceae, Sphingomonadaceae, Rhodobacteraceae, Bacillaceae, Hyphomicrobiaceae). Bacillales and reduced Alphaproteobacteria have a higher relative abundance at the GM site M4 compared to the rest of the AM sites

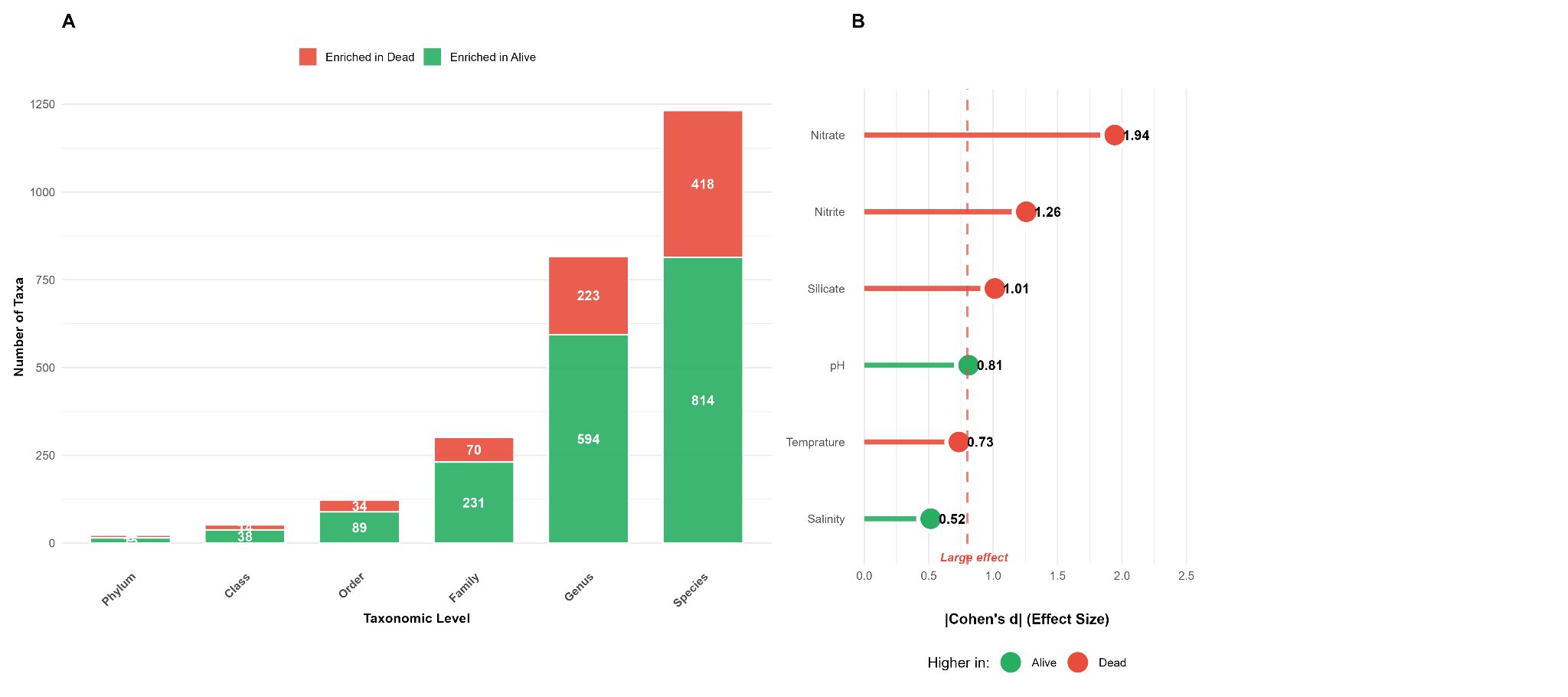

SFigure 5: Bacterial taxa and environmental parameters differentiating alive and dead mangrove sites.(A) Number of differentially enriched taxa per taxonomic level (selection criteria: ≥2 of contribution ≥1%, |Cohen's d| ≥0.5, |log₂FC| ≥0.5, Wilcoxon p<0.1). Green, enriched in alive magrove site; red, enriched in dead mangorve site. (B) Top ten environmental parameters by effect size. Horizontal lines indicate large (d=0.8, dashed) and medium (d=0.5, dotted) effect thresholds. Colors as in (A). Distance metrics: log-Jensen-Shannon (bacteria), Hellinger-Euclidean (parameters).

Additional Figure 5 illustrates the variation in enrichment and the associations of bacteria across ecosystem states. Following extremely high criteria (Cohen d > |.5|, Wilcoxon p < 0.1,log₂FC > |0.5|, ≥2% dissimilarity contribution) alive mangrove (AM) sites contained much more enriched taxa at all taxonomic levels: 65.2% of phyla (15/23), 73.1% of classes (38/52), 76.7% of families (231/301), 72.7% of genera (594/817). The Bacillales (d = 1.22, 20.0% contribution) and Alicyclobacillaceae (d = 0.39, 9.9%) were the key contributors of dissimilarity in dead mangrove (DM) sites (versus Hyphomicrobiales d = -1.54, 10.3%) and Hyphomicrobiaceae d = [?]1.91, 6.2%) in AM sites. Environmental analysis showed that there were strong geochemical effects (the highest effect of nitrate, d = 1.94, p = 0.027; 14.1-fold DM enrichment), then nitrite (d = 1.26), silicate (d = 1.01), pH decline (d = 0.81), and salinity reduction (d = -0.51). Nitrate, nitrite, and pH were verified as the principal discriminating axes by using the vector fitting (r > 0.80)..

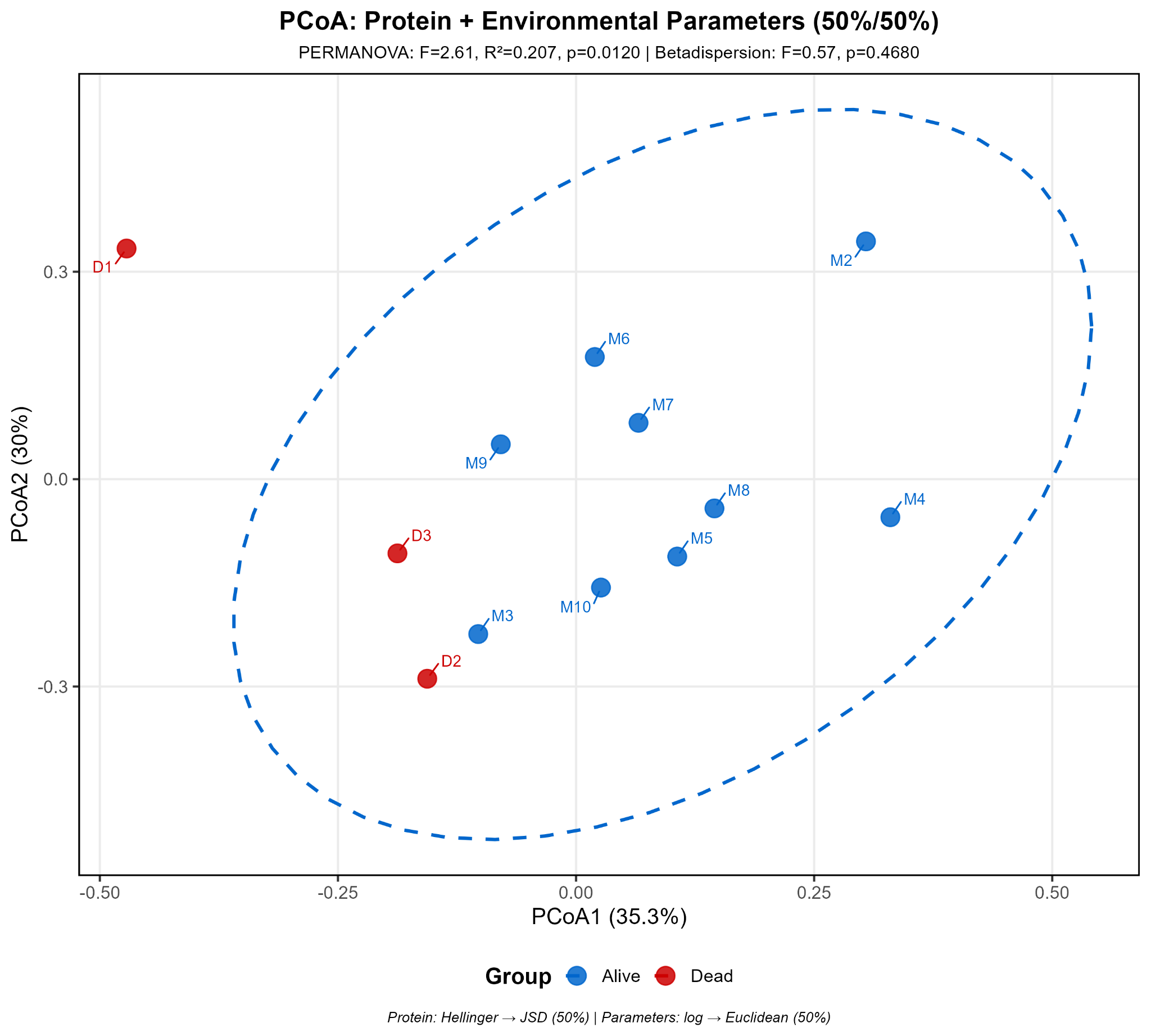

SFigure 6: Results of PCoA showing the metagenomic bacteria and PICRUSt2 KO data (Supplementary Material 4). PCoA of PICRUSt2 v2.5.3 predicted KO profiles (Jensen-Shannon distance, NSTI <2.0). PERMANOVA: F₁,₁₁ = 2.61, R² = 0.207, p = 0.0120 (significant). Betadisper: F = .57, p = 0.468 (homogeneous).

Principal Coordinates Analysis of predicted KEGG Orthology (KO) profiles (Jensen-Shannon distance; mean NSTI: 0.07 ± 0.02) is shown in Supplementary Figure 6; this analysis reveals significant functional divergence between alive mangrove (AM) and dead mangrove (DM) communities. PERMANOVA statistically confirmed the existence of substantial group separation (F = 2.61, R^2^ = 0.207, p = 0.012) and homogeneous dispersion (betadisper: F = 0.57, p = 0.468) was used to confirm that divergence is not due to differences in variance. The AM locations were oriented in ordination geometry to positive PC1, which is linked to biosynthetic and detoxification processes, and DM locations were oriented in ordination geometry to negative PC1/elevated PC2, which is related to stress-response activities. AM-enriched orthologs were phosphoribulokinase (K00799, 12.6-fold), NADPH dehydrogenase (K00059, 3.6-fold), ABC transporters (K01990, K01992, K02035; 2.5-3.7-fold) and DNA repair genes (K01999, K02004; 3.3-6.4-fold). The regulators of stress responses (sigma-70, K03088) were enriched in DM sites. The environment-related vector modelling correlated high nitrate/nitrite levels with DM-related stress and anaerobic metabolism genes, while the neutral pH was consistent with AM-related biosynthetic and detoxification capacity. The deterministic impact of functional restructuring in pollution stress is emphasised by confidence ellipses (85-95%) of the stress.
